# Antibiotic-degrading resistance changes bacterial community structure via species-specific responses

**DOI:** 10.1101/2022.12.21.521377

**Authors:** Ayush Pathak, Daniel C. Angst, Ricardo León-Sampedro, Alex R. Hall

**Affiliations:** Institute of Integrative Biology, Department of Environmental Systems Science (D-USYS), ETH Zurich, Zurich, Switzerland

**Keywords:** Exposure protection, antibiotic detoxification, beta-lactamases, OXA-48, piperacillin-tazobactam

## Abstract

Some bacterial resistance mechanisms degrade antibiotics, potentially protecting neighbouring susceptible cells from antibiotic exposure. We do not yet understand how such effects influence bacterial communities of more than two species, which are typical in nature. Here, we used experimental multispecies communities to test the effects of clinically important pOXA-48-plasmid-encoded resistance on community-level responses to antibiotics. We found resistance in one community member reduced antibiotic inhibition of other species, but some benefitted more than others. Further experiments with supernatants and pure-culture growth assays showed the susceptible species profiting most from detoxification were those that grew best at degraded antibiotic concentrations (greater than zero, but lower than the starting concentration). This pattern was also observed on agar surfaces. By contrast, we found no evidence of a role for higher-order interactions or horizontal plasmid transfer in community-level responses to detoxification in our experimental communities. Our findings suggest carriage of an antibiotic-degrading resistance mechanism by one species can drastically alter community-level responses to antibiotics, and the identities of the species that profit most from antibiotic detoxification are predicted by their intrinsic ability to grow at degraded antibiotic concentrations.

## Introduction

Antibiotic resistance is an obstacle to effective treatment of bacterial infections (1,2). Individual strains or species carrying resistance mechanisms often coexist with other species in multispecies communities. Therefore, to predict responses to antibiotics at the level of individual species or entire communities/microbiomes, we need to understand how resistance mechanisms carried by one species affect neighbouring species in the same community (3,4). One common mechanism by which some resistance mechanisms can influence surrounding microbes is by reducing the antibiotic concentration, which can in turn benefit neighbouring susceptible cells (5), sometimes called exposure protection (3,6). Important examples of this type of detoxification include carbapenemases and other beta lactamases (7–11), which enzymatically deactivate beta-lactam antibiotics. Past work showed exposure protection can occur between strains of the same species or pairs of species (3,6–11). This suggests detoxification is an important mechanism by which resistance can influence community structure. However, it is not yet known whether or how detoxifying resistance mechanisms influence community structure (relative abundances of different species) in communities of more than two species.

There are multiple ways community structure could change in response to antibiotic degradation. For example, if resident species vary in their ability to grow at degraded antibiotic concentrations, the identities of the species that profit most from detoxification may be predictable from their abilities to grow in pure cultures at relevant concentrations. Depending on the extent of degradation, this may in turn be correlated with interspecific variation of widely measured parameters such as the minimum inhibitory concentration (MIC) or growth rate in the absence of antibiotics (12). Alternatively, if some of the resident species can become resistant to the extant antibiotic concentration, either via horizontal transfer or other mechanisms, community structure upon detoxification may instead reflect variable rates of resistance evolution. A third possibility is that community-level responses to antibiotic degradation depend on higher order interactions, such as growth of some susceptible species in the presence of the resistant species being influenced by interactions with other susceptible species. There is some evidence that higher-order interactions can occur among microbes (13,14), but as of yet there is no consensus about their importance for microbial community structure upon antibiotic exposure (3,15–17). Thus, understanding which resident, susceptible species profit most (increase most in relative abundance) when a resistant species detoxifies the local environment would improve our basic understanding of how antibiotic resistance influences microbial diversity, and our ability to predict community-level responses to antibiotics in nature or in treatment contexts.

To test how a potentially detoxifying resistance mechanism affects the relative abundances of different species, we assembled experimental, multi-species communities comprising *Escherichia coli, Staphylococcus aureus, Salmonella enterica* serovar Typhimurium, *Enterococcus faecalis, Pseudomonas aeruginosa* and *Klebsiella pneumoniae*. We chose these species because they are phylogenetically and phenotypically diverse, and are all associated with humans, either as commensals (18,19) or as pathogens (20–23). We tested whether community structure was affected by carriage of a clinically relevant plasmid encoding a carbapenemase, pOXA-48 (24–26), by one community member (*E. coli*). We monitored plasmid effects on community structure both with and without antibiotics, using a clinically relevant combination of piperacillin and tazobactam (27). Piperacillin is a piperazine derivative of ampicillin (28,29), generally used in combination with the beta-lactamase inhibitor tazobactam, a penicillanic acid sulfone derivative (30,31). We found plasmid carriage by *E. coli* affected community structure via antibiotic detoxification, and the species benefitting most from this were identified by their relative abilities to grow at degraded antibiotic concentrations.

## Materials and methods

### Bacteria and growth media

We used seven strains (from six species; Table S1) to assemble multispecies communities: *Escherichia coli* K-12 MG1655 (with and without pOXA-48 plasmid), *Staphylococcus aureus* subsp. *aureus* (Type Strain), *Salmonella enterica* serovar Typhimurium SL1344, *Enterococcus faecalis* JH2-2, *Pseudomonas aer*μ*ginosa* PAO1, *Klebsiella pneumoniae* subsp. *pneumoniae* (Schroeter) Trevisan (Type Strain). We cultured bacteria in lysogeny broth (LB) media or on Liofilchem Chromatic™ MH agar (Roseto degli Abruzzi, Italy), incubated at 37°C for 24 hours. Each of the six species form colonies with distinct colour/morphology on chromatic agar, allowing us to estimate community composition and total abundance by plating.

### Plasmid and antibiotics

We used a 63.6kB pOXA-48-like IncL broad-host range conjugative plasmid carrying a single beta-lactamase gene *bla*_OXA-48_ which confers resistance to most beta-lactam antibiotics (25). This plasmid was obtained from a clinical *Escherichia coli* strain isolated in the Division of Clinical Microbiology at the University Hospital Basel, Switzerland (32). Specifically, this pOXA-48-like plasmid (acc. num. UWXP01000003.1) shares a 97% coverage and >99% identity both with the first described pOXA-48 plasmid (33) and with pOXA48_K8, one of the best studied pOXA48-like variants (34). To test whether the pOXA-48 plasmid used in this study was conjugative, we used a mating assay between the native clinical strain carrying the plasmid and a chloramphenicol resistant *E. coli* K-12 MG1655 (CmR, Δ*galK::cat*), both on agar and in liquid (Figure S1). In our main experiments described below, we assembled communities with and without the pOXA-48 plasmid by including plasmid-carrying or plasmid-free versions of *E. coli* K-12 MG1655. For communities treated with antibiotics, we used a combination of 7.5 μg/ml piperacillin and 1.5 μg/ml tazobactam, unless stated otherwise. In some experiments we used selective chromatic agar plates, including either 7.5 μg/ml or 10 μg/ml piperacillin and 1.5 μg/ml or 2 μg/ml tazobactam.

### Assembling experimental communities with/without antibiotics and plasmid

To test how the structure of the community was affected by antibiotics and the plasmid, we cultured five replicated communities in four treatments (all combinations of +/- antibiotic and +/- plasmid). For each replicate microcosm, we first cultivated all constituent species in separate overnight cultures, each inoculated from separate colonies, in 200 μl LB in 96-well microplates. We used independent sets of overnight pure cultures (one per constituent species) for every replicate microcosm. For inoculation of a given microcosm, we diluted the overnight culture of each constituent species 1000-fold into a total volume of 200 μl. Before and after 24h incubation, we plated dilutions from each microcosm on chromatic agar to estimate initial and final abundance and community composition. To test for the emergence of antibiotic resistance in susceptible species incubated with the plasmid-carrying strain (for example via conjugative plasmid transfer), we picked all colonies (37 *S*. typhimurium and 13 *P. aeruginosa*) that survived in any of the five replicate microcosms of the antibiotic+plasmid treatment (except for the plasmid-carrying *E. coli*). We then restreaked these colonies on chromatic agar with piperacillin and tazobactam to check for stable growth at concentrations inhibitory to ancestral strains. As a second test for resistant variants, we plated 20 μl aliquots from all microcosms in all treatments directly onto chromatic agar with piperacillin and tazobactam.

### Supernatant experiments

We tested whether communities incubated with vs without the plasmid (with antibiotics) modified the abiotic environment in different ways, and whether this had variable downstream effects on growth of different constituent species. To do this, we first produced three types of supernatant, by incubating communities as in the main experiment in the antibiotic+plasmid treatment, in the antibiotic+no plasmid treatment, and as a control by incubating sterile medium with antibiotics. We incubated 96 replicate microcosms as above in each treatment, before pooling the replicates in each treatment (to obtain sufficient culture volume) and filtering (0.45 μm), producing 10 ml total of supernatant per treatment, which we froze at -20 °C. We then tested whether all seven strains could grow in pure culture at various concentrations of each supernatant type, diluted in fresh LB. To compensate for the effect of nutrient depletion during supernatant preparation, which could potentially create a false signal of growth inhibition in downstream experiments, we added 10% and 5% of concentrated LB solution (at four times the standard concentration) to treatments with undiluted and 50% supernatant, respectively. This makes interpretation of quantitative growth scores in these treatments relative to other concentrations problematic, but enables us to rule out nutrient depletion as an explanation for zero growth of some species in these supernatants. Supernatant cultures were inoculated in 100 μl microcosms from overnight cultures by 1000-fold dilution as above, with three replicates in each combination. We estimated growth after 24 hours by Optical Density (600 nm) with an Infinite M200 Pro NanoQuant Tecan plate reader (Männedorf, Switzerland).

### Testing for species-specific benefits of antibiotic detoxification on agar

After finding evidence of species-specific effects of detoxification in liquid culture, we made a second test on solid media, where some past work has shown evidence of exposure protection between pairs of strains (8). First, we streaked plasmid-carrying *E. coli* across the diameter of 18 chromatic agar plates supplemented with 7.5 μg/ml piperacillin and 1.5 μg/ml tazobactam. We then streaked each antibiotic-susceptible strain perpendicularly to the plasmid-carrying *E. coli*, with three plates per species. We left ∼0.5 cm between the strains on each plate. We then incubated all plates and checked for evidence growth of the susceptible strain was increased by proximity to the plasmid-carrying strain. To test for such interactions in communities of more than two species, we plated diluted cultures of assembled communities of susceptible species onto rectangular plates with a left-to-right antibiotic gradient and a streak of plasmid-carrying *E. coli*. To prepare gradient plates, prior to adding bacteria, we poured and dried a slanted slab of 20 ml LB agar containing 5 µg/ml piperacillin and 1 µg/ml tazobactam, before adding a further 20 ml antibiotic-free LB agar on top; we dried plates before adding bacteria.

### Antibiotic dose-response curves and species-specific growth without antibiotics

To estimate the intrinsic ability of each species to grow at relevant antibiotic concentrations, we tested susceptibility to various concentrations of piperacillin+tazobactam. We prepared each culture as above, inoculating three replicates per strain via 1000-fold dilution, each from an independent overnight culture. We used a two-fold dilution series of piperacillin in LB, ranging from 37.5 to 0.585 μg/ml, plus 0 μg/ml, supplemented with 15 μg/ ml of tazobactam at all concentrations of piperacillin except 0 μg/ml. After 24 hours, we measured optical density (600 nm) as above. The MIC for each strain was the concentration at which there was no detectable growth, using a cut-off for detectable growth of OD (600 nm) ≥ 0.05. In a separate experiment, we also measured the growth rate of each species in pure culture without antibiotics. These cultures were inoculated as above in LB, and incubated in the plate reader, measuring optical density (600 nm) every 15 minutes for 24 hours.

### Testing for changes in community structure at artificially reduced antibiotic concentrations

We hypothesized artificially reducing the antibiotic concentration would result in similar community composition in microcosms without plasmid-carrying *E. coli* compared to communities from the antibiotic+plasmid treatment of the main experiment. That is, we aimed to mimic the detoxifying effect of the plasmid by extraneously reducing the antibiotic concentration. We inoculated four replicate communities as above, without the plasmid-carrying strain, at various antibiotic concentrations (7.5 μg/ml piperacillin and 1.5 μg/ml tazobactam, 0.9 μg/ml piperacillin and 0.18 μg/ml tazobactam, 0.46 μg/ml piperacillin and 0.09 μg/ml tazobactam, 0.23 μg/ml piperacillin and 0.04 μg/ml tazobactam and 0 μg/ml of piperacillin and tazobactam, equivalent to 100%, 12.5%, 6.25%, 3.125% and 0% of the concentration used in the main experiment). After incubation, we inferred community structure and abundance by plating as above.

### Pairwise interactions between plasmid-carrying *E. coli* and other species

We tested whether species benefiting most from detoxification in the main experiment above also benefited most from detoxification in two-species cultures with only the plasmid-carrying *E. coli*. In other words, we asked whether the benefits of detoxification were specific to six-species communities (e.g., because they rely on higher-order interactions), or if they could be explained by pairwise interactions between individual susceptible species and the plasmid-carrying *E. coli* strain. We cultivated each species in pure cultures (without antibiotics), and in cocultures with plasmid-carrying *E. coli* (both with and without antibiotics). We prepared cultures as above, inoculating by 1000-fold dilution from overnight cultures of each species (that is, the total number of cells was higher in the two-species cultures than in the pure cultures). This set-up tests whether growth of each species is altered by addition of plasmid-carrying *E. coli* relative to how it grows on its own when inoculated at the same density, following past work (35,36). We then incubated and plated as above, also including a separate plating on piperacillin+tazobactam to test for resistance evolution in initially susceptible species.

### Statistical Analysis

All statistical analyses were conducted in R version 4.1.0. To analyse community structure, we used principal component analysis of the absolute abundances (CFU/ml) of each species in different treatments of the main experiment. The variables were shifted to be zero centered and were scaled to have unit variance. We also used permutational multivariate analysis of variance (permANOVA) based on Bray-Curtis dissimilarity matrix, using the “vegan” R package (37,38). To estimate growth rates in pure culture without antibiotics, we used the “fitR” package (39). This package uses a sliding-window approach to estimate the maximum slope of OD over time for a defined number of consecutive points (here, five consecutive time points with intervals of 15 minutes).

## Results

### Resistant *E. coli* rescues *P. aeruginosa* and *S*. Typhimurium from antibiotic inhibition

When we cultivated experimental multispecies communities with antibiotics, inclusion of a resistance plasmid in *E. coli* reduced the effect of antibiotics on total bacterial abundance (Figure 1A, two-way ANOVA, plasmid × antibiotic interaction: F_1,16_=229.5, p<0.001). Community structure, in terms of relative abundances of different species (Figure 1B-C), was also significantly affected by both antibiotics and by the plasmid (permANOVA, effect of plasmid: pseudo-F_1,16_=14.534, p=0.001; effect of antibiotic: pseudo-F_1,16_=16.955, p=0.001). In the absence of the plasmid, antibiotic treatment resulted in high variability among replicate microcosms, and fewer species being detected at the end of the experiment (Figure 1C). Total bacterial abundance in these microcosms (with the antibiotic, without the plasmid) was lower after 24h than at 0h, indicating community structure here reflected a small fraction of surviving cells from some species, rather than variable population growth across the plasmid-free species (mean total abundance after 24 hours = 95 CFU/ml, SD=106.65, mean at 0 hours = 2.2 × 10^6^ CFU/ml, SD=1.3 × 10^6^). The plasmid had little effect on community composition in the absence of antibiotics, but upon antibiotic treatment it enabled *P. aeruginosa, S*. Typhimurium and *E. coli* to grow (antibiotic × plasmid interaction in permANOVA: pseudo-F_1,16_=14.694, p=0.001; Figure 1C). Thus, as well as increasing growth of the strain carrying the plasmid (*E. coli*) in the presence of antibiotics, the plasmid rescued some otherwise susceptible community members (*P. aeruginosa* and *S*. Typhimurium) from antibiotic inhibition.

**Figure 1.**
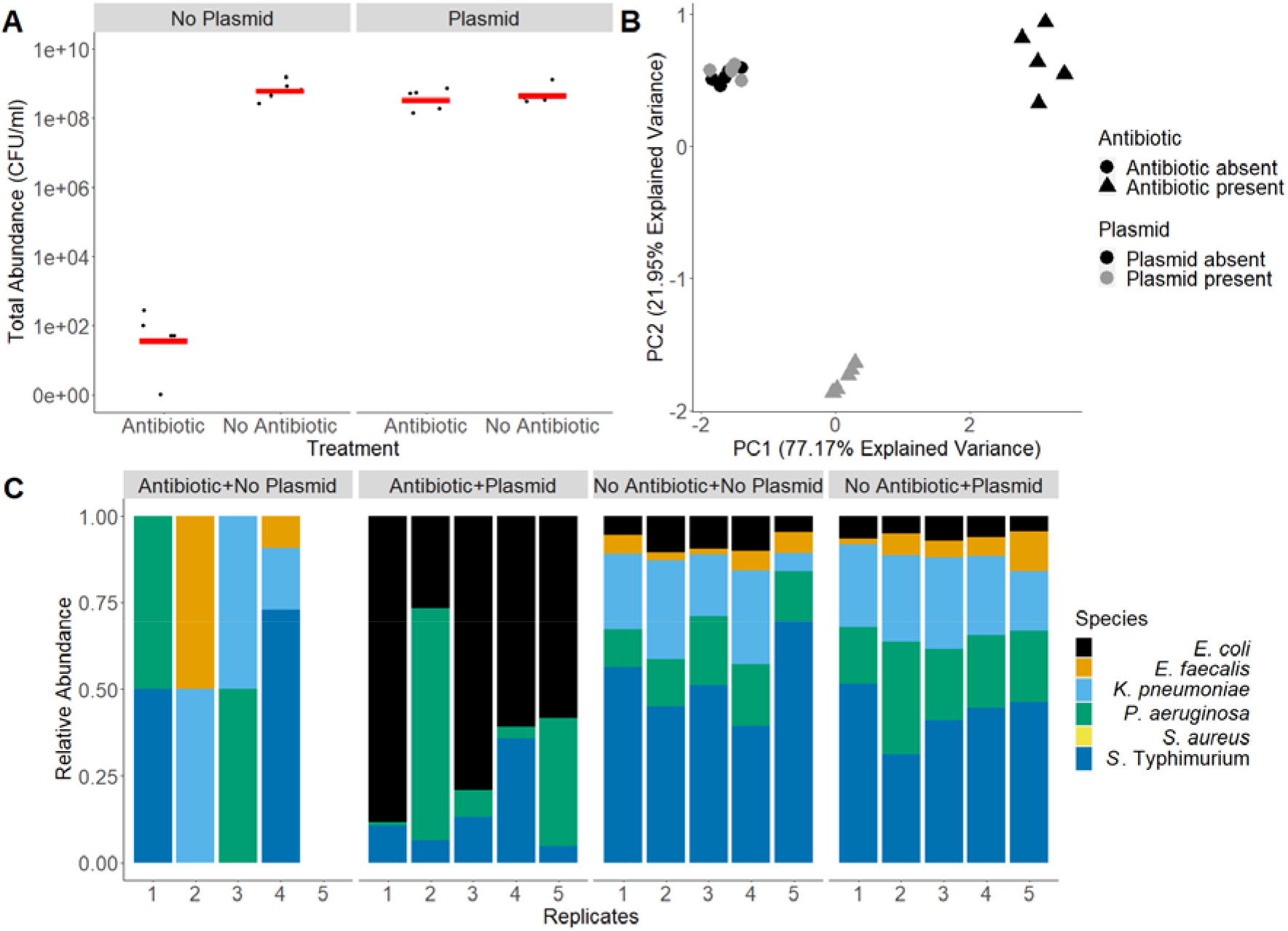
Community abundance and structure with/without antibiotics and a resistance plasmid. A. Estimated total abundance (CFU/ml) in five replicated microcosms in each treatment (with/without a resistance plasmid carried by *E. coli*, and with/without antibiotic treatment with piperacillin+tazobactam). B. Principal Component Analysis of community structure in the different treatments; input variables are absolute abundances of the six species in each microcosm. C. Community structure in different treatments; estimated relative abundances of each species in five replicate microcosms in each treatment.

### No evidence of resistance evolution in antibiotic-susceptible species

Variation among the susceptible species in terms of their abilities to acquire antibiotic resistance during the experiment (e.g., by horizontal transfer from the plasmid-carrying strain) is a potential explanation for why some species performed better than others in the presence of plasmid-carrying *E. coli* plus antibiotics. We tested this by restreaking colonies from each of the initially susceptible species from the end of the main experiment onto antibiotic agar. We picked all *S*. Typhimurium and *P. aeruginosa* colonies (37 *S*. Typhimurium colonies and 13 *P. aeruginosa* colonies) from five chromatic agar plates from the antibiotic+plasmid treatment, and restreaked them on chromatic agar supplemented with antibiotics. None of the re-streaked colonies formed new colonies on antibiotic agar. As a second test, we plated 20 μl aliquots from all microcosms of the four treatments (Figure 1) on chromatic agar plates with 10 μg/ml piperacillin and 2 μg/ml tazobactam. On these plates the only colonies we observed were of *E. coli* with the pOXA-48 plasmid. Thus, we found no evidence that the rescue of susceptible species caused by addition of the plasmid in the presence of antibiotics (observed above) was linked to acquisition of resistance during the experiment. Despite this, a separate mating experiment between the native clinical donor strain and *E. coli* MG1655_CmR showed the pOXA-48 plasmid was conjugative at high rates in other experimental conditions (Figure S1), as previously reported for other variants (34).

### Plasmid carriage by *E. coli* detoxifies the environment for other species

We next tested whether plasmid carriage by *E. coli* modified the abiotic environment in the microcosms compared to the same communities without plasmids, by culturing each species individually in three types of supernatants: (i) supernatant from communities incubated as in the main experiment in the antibiotic + plasmid treatment, (ii) from communities incubated as in the antibiotic + no plasmid treatment, and (iii) sterile medium with antibiotics, as a control (Figure 2). We found for all susceptible species (species without the plasmid), population growth was higher in supernatants from communities with the plasmid than without (pink and yellow series in Figure 2). This suggests the plasmid reduced the antibiotic concentration in the medium. The magnitude of this effect varied among the susceptible species (two-way ANOVA with growth in undiluted supernatant as response variable, excluding control supernatant, species × supernatant type interaction: F_5,24_= 26.43, p<0.01). *P. aeruginosa, K. pneumoniae, S. aureus* and *S*. Typhimurium all showed relatively strong responses to detoxification (difference between plasmid+ and plasmid-supernatant types) compared to *E. faecalis* and plasmid-free *E. coli*. For *P. aeruginosa* and *S*. Typhimurium, this resulted in population densities at the highest supernatant concentration close to those observed in supernatant-free LB (Figure 2). Both types of community-derived supernatant (with the plasmid and without) supported more growth than control supernatant without bacteria, indicating there was some antibiotic detoxification even without the plasmid (Figure 2). In summary, communities with the plasmid rendered the abiotic environment less growth-inhibitory, but some species were better than others at exploiting this.

**Figure 2.**
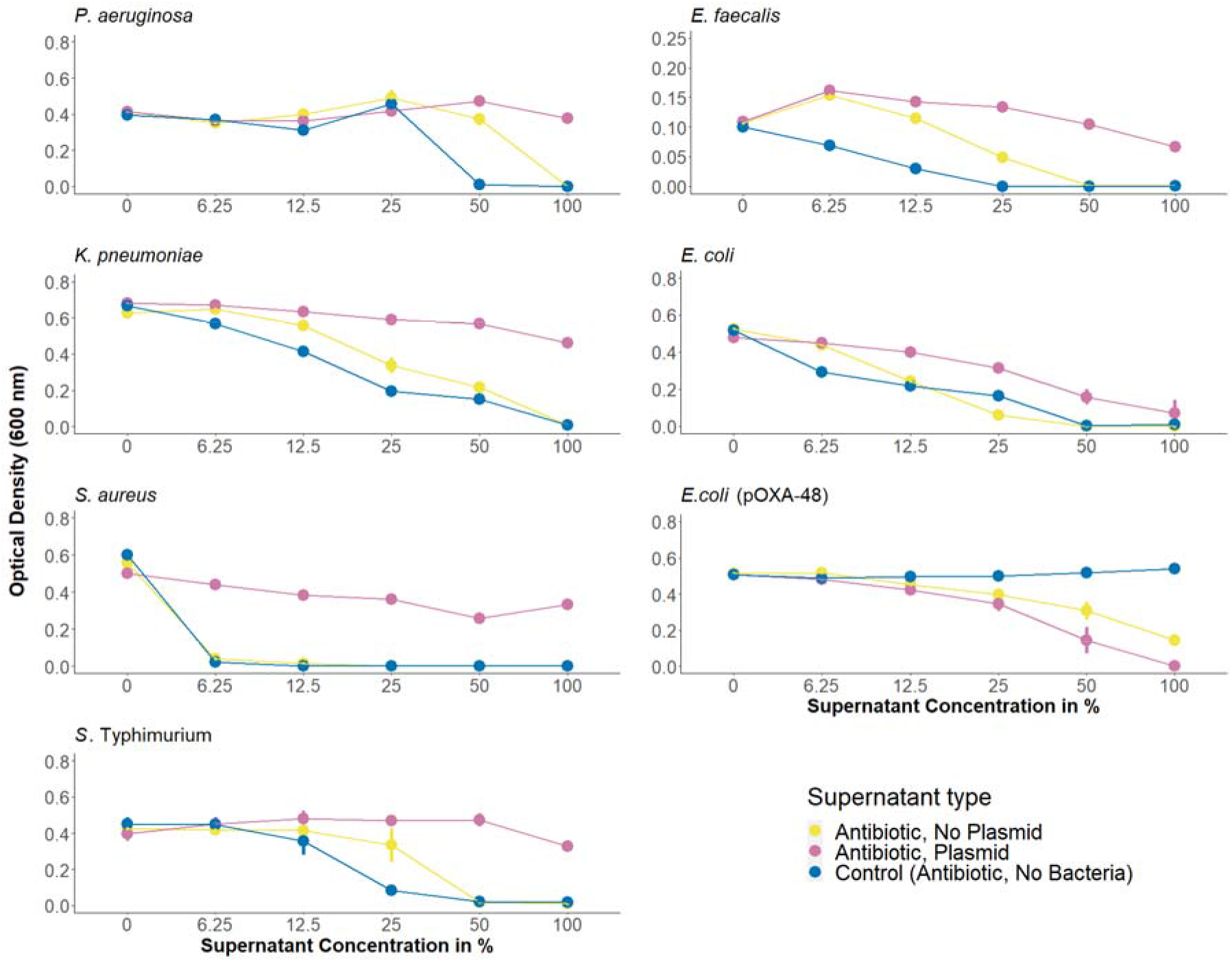
Abundance (optical density at 600 nm) of each species after 24h growth at various concentrations of three types of supernatant. Supernatants were extracted from community cultures with and without the plasmid, or sterile medium, incubated with antibiotics. Each species was then inoculated into various concentrations of supernatant (*x*-axis, given as the percentage by volume of supernatant diluted in fresh LB medium; note at 50% and 100%, a small volume of concentrated LB media was added to account for nutrient depletion, see methods). Note the different *y*-axis scale for *E. faecalis*, used because of the much lower data range for this species.

### Species-specific benefits of antibiotic degradation on agar

As a second test for species-specific benefits of detoxification by the plasmid-carrying strain, we streaked all antibiotic-susceptible species from the antibiotic+plasmid treatment of the main experiment perpendicularly to the plasmid-carrying *E. coli* on chromatic agar with antibiotics (Figure 3A). After incubation, only *P. aeruginosa* and *S*. Typhimurium showed visible growth, and only in proximity to the plasmid-carrying strain. In a further test for interactions on agar, but at the community level, we plated entire multispecies communities on agar with a gradient of piperacillin+tazobactam, with the plasmid-carrying *E*. coli streaked across the middle (Figure 3B). This showed again that *P. aeruginosa* and *S*. Typhimurium were among the species best able to exploit detoxification (colonies on the high-antibiotic side of the plate in proximity to the plasmid-carrying strain; Figure 3B & S4). This supports the species-specific benefits of detoxification we observed above in liquid culture, and shows this can also apply in a spatially structured environment.

**Figure 3.**
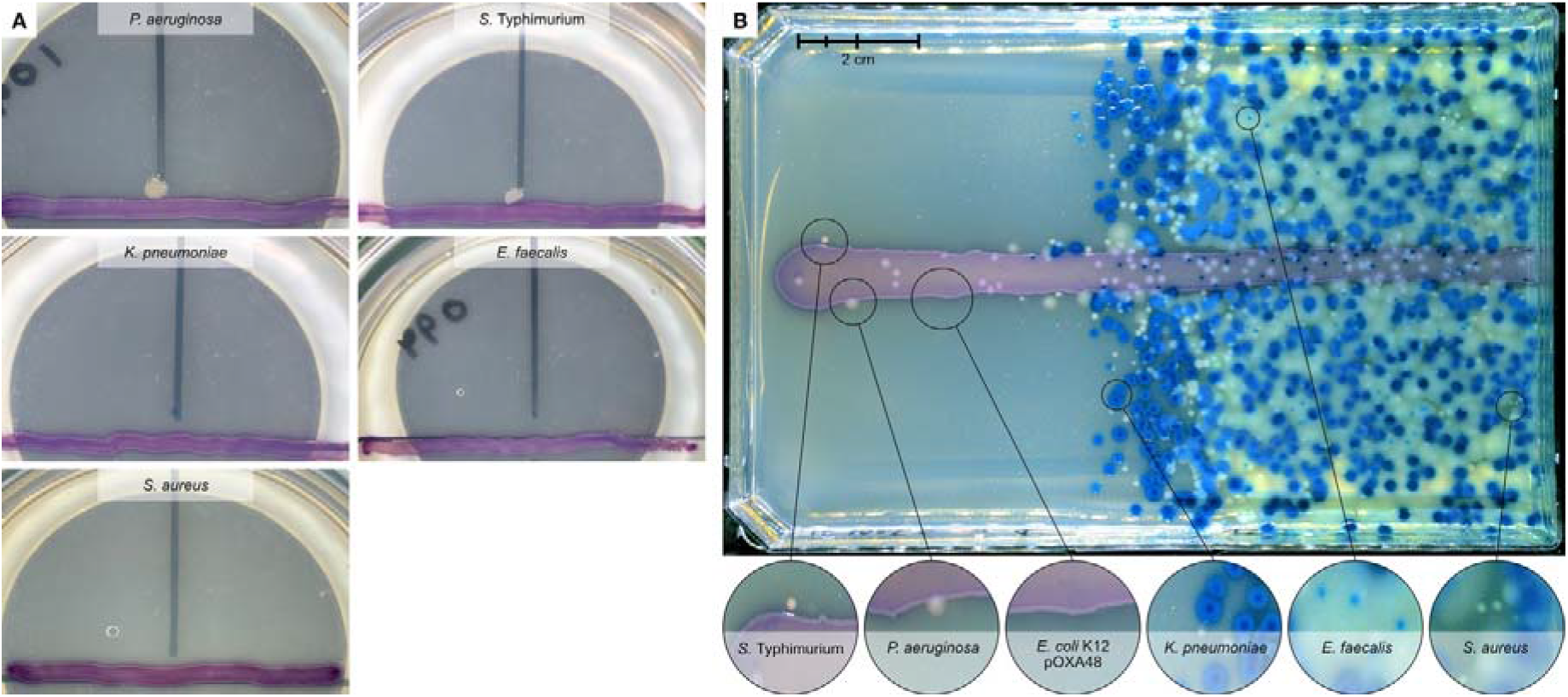
Species-specific benefits of detoxification on agar. (A) Two out of five antibiotic-susceptible species (*P. aeruginosa* and *S*. Typhimurium) show visible growth on antibiotic-agar in proximity to a resistant, plasmid-carrying *E. coli* strain. Each susceptible species was streaked on chromatic agar with antibiotics (vertical black line) perpendicular to the pOXA-48 carrying *E. coli* (purple horizontal line) and incubated for 24 hours. Each combination was replicated three times and one replicate is shown; other replicates were similar in all combinations (three out of three for *P. aeruginosa* and two out of three for *S*. Typhimurium were qualitatively identical; see Figure S3. (B) Multispecies communities plated on a gradient of piperacillin+tazobactam (left-to-right; maximum concentration in overlay agar =5 µg/ml piperacillin and 1 µg/ml tazobactam) with the plasmid-carrying *E. coli* strain streaked horizontally (Figure S4).

### Species that profit most from detoxification also grow relatively well at degraded antibiotic concentrations in pure culture

One possible explanation for species-specific benefits of detoxification is that some species grow better than others at degraded antibiotic concentrations. The relevant range of concentrations in our experiments includes 7.5 μg/ml (the starting concentration used in the main experiment) and non-zero concentrations lower than 7.5 μg/ml (our supernatant assay above showed communities with the plasmid reduce antibiotic inhibition, but do not remove it entirely). In pure culture, no species except the plasmid-carrying *E. coli* showed significant growth at 7.5 μg/ml piperacillin or higher (Figure 3). At non-zero concentrations lower than 7.5 μg/ml, *P. aeruginosa* and *S*. Typhimurium were the best-performing susceptible species at all but one of the tested concentrations (Figure 4). Thus, the species that profited most from antibiotic detoxification by plasmid-carrying *E. coli* in our main experiment (Figure 1), in supernatant assays (Figure 2), and on agar-plate assays (Figure 3), were those that grew best in pure cultures at degraded antibiotic concentrations. Note growth at degraded concentrations was not simply predicted by the relative MICs of susceptible species: *Klebsiella pneumoniae* (MIC: 9.375 μg/ml) had a higher MIC than *P. aeruginosa* (MIC: 4.65 μg/ml) and *S*. Typhimurium (MIC: 4.65 μg/ml) but performed worse at most non-zero concentrations lower than 7.5 μg/ml. The relatively high MIC of *K. pneumoniae* also agrees with its dense growth at higher antibiotic concentations compared to other species on agar in Figure 3B.

**Figure 4.**
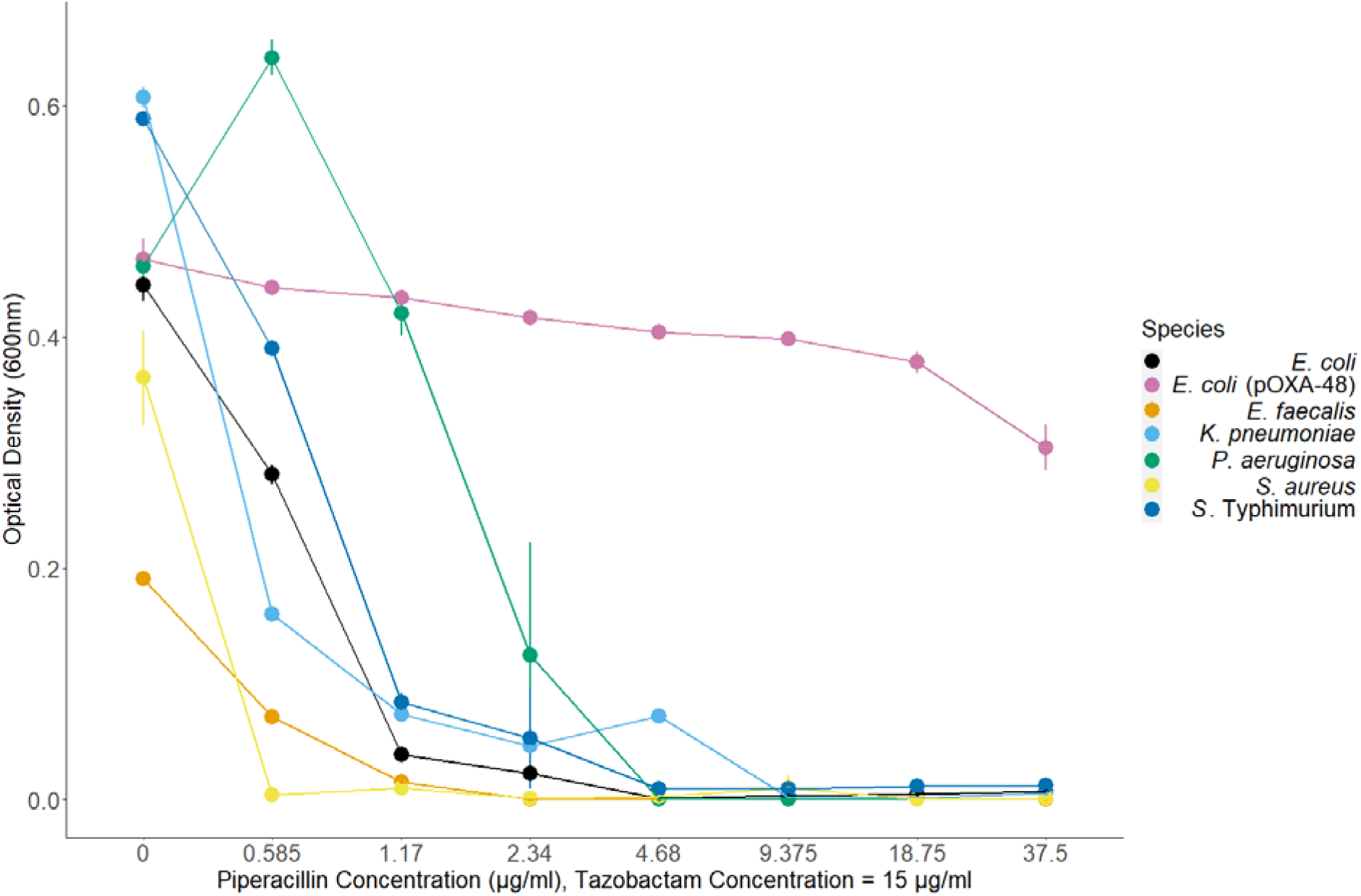
Dose-response curves for all species with varying concentrations of piperacillin and a constant tazobactam concentration. Error bars represent 95% confidence intervals (*n* = 3).

Unlike growth at degraded antibiotic concentrations, the relative abilities of different species to grow in the absence of antibiotics did not explain the success of *P. aeruginosa* and *S*. Typhimurium in the antibiotic+plasmid treatment of the main experiment. This is shown firstly by another species (*K. pneumoniae*) reaching abundances similar to or greater than *P. aeruginosa* in plasmid-free, antibiotic-free microcosms of the main experiment, but being excluded in the antibiotic+plasmid treatment (Figure 1). It is also possible that relative final abundances in community microcosms with antibiotic and the plasmid depend more on the intrinsic growth rates of individual species during exponential phase without antibiotics, rather than on their relative final abundances without antibiotics as measured above (Figure 1). To investigate this, we estimated the maximum growth rate in the absence of antibiotics for each species (Figure S2). This showed that, for both *P. aeruginosa* and *S*. Typhimurium, there was at least one other antibiotic-susceptible species with a similar or higher maximum growth rate in the absence of antibiotics (Figure S2; effect of species in one-way ANOVA: p<0.05; *S. aureus* and *S*. Typhimurium > *P. aeruginosa*; *S. aureus, S*. Typhimurium and *K. pneumoniae* > *E. faecalis* by Tukey’s HSD comparisons). Final abundances in the same pure culture assays showed that neither *P. aeruginosa* nor *S*. Typhimurium had significantly higher abundances than other species (Figure S2).

### Artificially reducing the antibiotic concentration reproduces some of the same dynamics as observed upon plasmid introduction

We hypothesized our results were driven by plasmid-encoded resistance detoxifying the local abiotic environment. If this is true, artificially reducing the concentration of antibiotic, even in the absence of plasmid-encoded resistance, should result in some of the same changes in community structure as observed in the antibiotic + plasmid treatment of the main experiment. Without antibiotics, as expected, community structure resembled that of the antibiotic-free communities from our main experiment (Figure 5, 0%). At intermediate concentrations (greater than zero but lower than that used in the main experiment), *P. aeruginosa* was the most abundant susceptible species. *S*. Typhimurium also performed well in some microcosms at all intermediate concentrations tested. This supports our other results indicating these two species profit most from plasmid-mediated detoxification. We nevertheless interpret this experiment with caution, because changing the starting concentration does not recapitulate all possible effects from the antibiotic + plasmid treatment of our main experiment, such as the temporal dynamics of antibiotic degradation or the effect of the plasmid on growth of its *E. coli* host. Therefore, while this experiment is consistent with key effects of the plasmid arising from detoxification alone, we do not rule out other effects.

**Figure 5.**
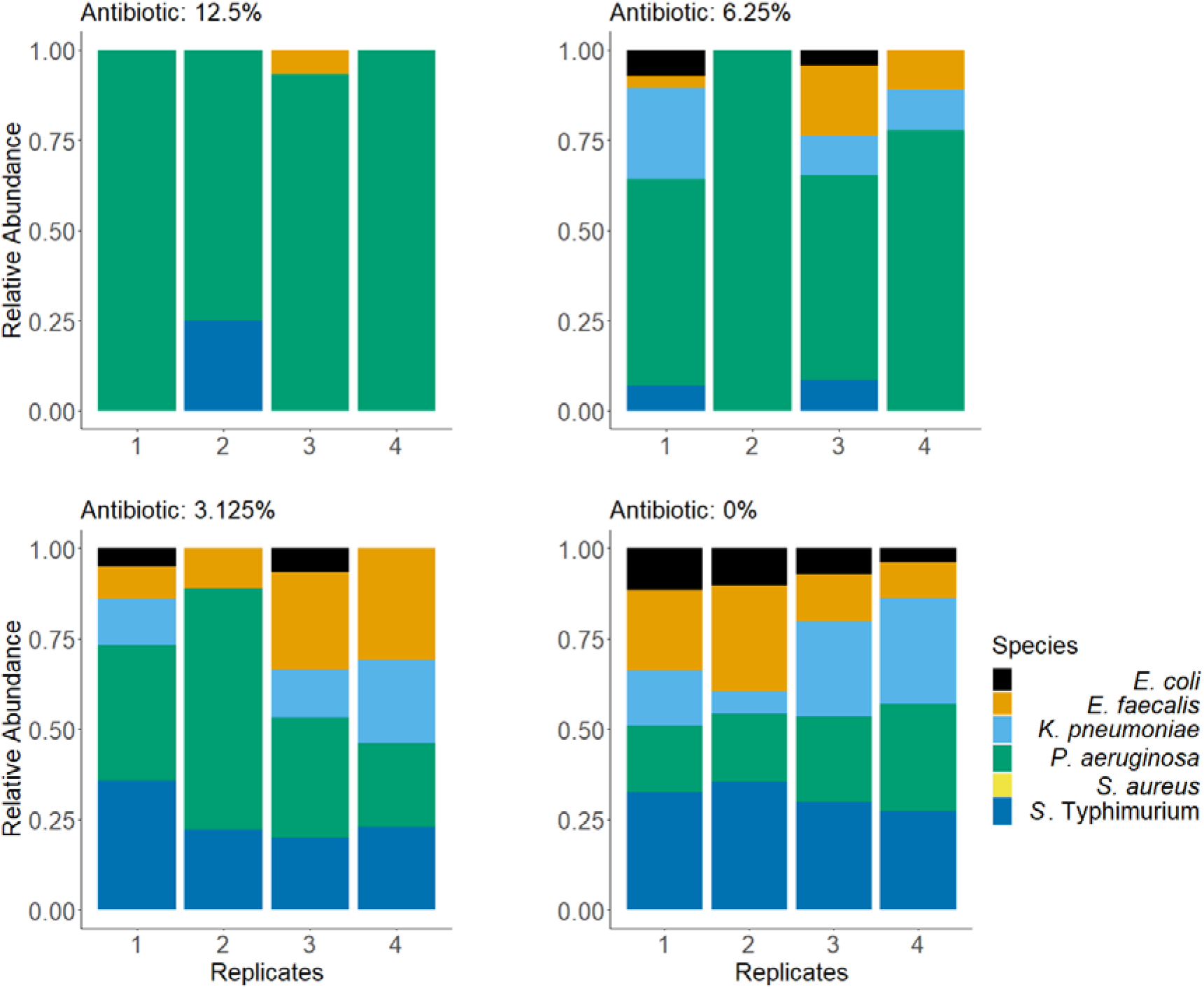
Community structure in microcosms without plasmid-carrying *E. coli* at various starting concentrations of antibiotics. Each panel shows a different antibiotic concentration given as a percentage of the starting concentration in the main experiment (Figure 1): 12.5% = 0.9 μg/ml piperacillin and 0.18 μg/ml tazobactam, 6.25% = 0.46 μg/ml piperacillin and 0.09 μg/ml tazobactam, 3.125% = 0.23 μg/ml piperacillin and 0.04 μg/ml tazobactam, 0% = No antibiotic. The 100% treatment (7.5 μg/ml piperacillin, 0.18 μg/ml tazobactam) is not shown; we detected negligible growth in this treatment, with 3 out of 4 replicates having no colonies at all.

### Higher-order interactions are not required to explain species-specific benefits of detoxification

The above experiments suggest *P. aeruginosa* and *S*. Typhimurium profit most from detoxification because they grow better than other species at degraded antibiotic concentrations, and therefore higher-order interactions are not required to explain changes in community structure upon detoxification. In this case, we would expect to see similar patterns as in the main experiment when each susceptible species is incubated only with the plasmid-carrying *E. coli* strain (without other susceptible species). We found this to be the case: in co-cultures with the plasmid-carrying strain plus antibiotics, only *S*. Typhimurium and *P. aeruginosa* reached detectable population densities (Figure 6). As in our main experiment, plating on antibiotic agar revealed no evidence of resistant variants of susceptible species. In the same experiment we included treatments without antibiotics, both in pure and co-cultures. In both these treatments, some of the other species performed similarly well compared to *P. aeruginosa* and *S*. Typhimurium (Figure 6). There was variation among species here (p<0.05 by one-way ANOVA for both treatments; Figure 6), but this primarily reflected the poor growth of *S. aureus*. The other four species did not differ from each other significantly (Tukey’s HSD, p>0.05 for all pairwise comparisons among these four species in both treatments without antibiotics). This experiment therefore indicates, as above, *P. aeruginosa* and *S*. Typhimurium profited most from detoxification, even when the other susceptible species were absent, and this was not associated with superior growth of these two species in the absence of antibiotics.

**Figure 6.**
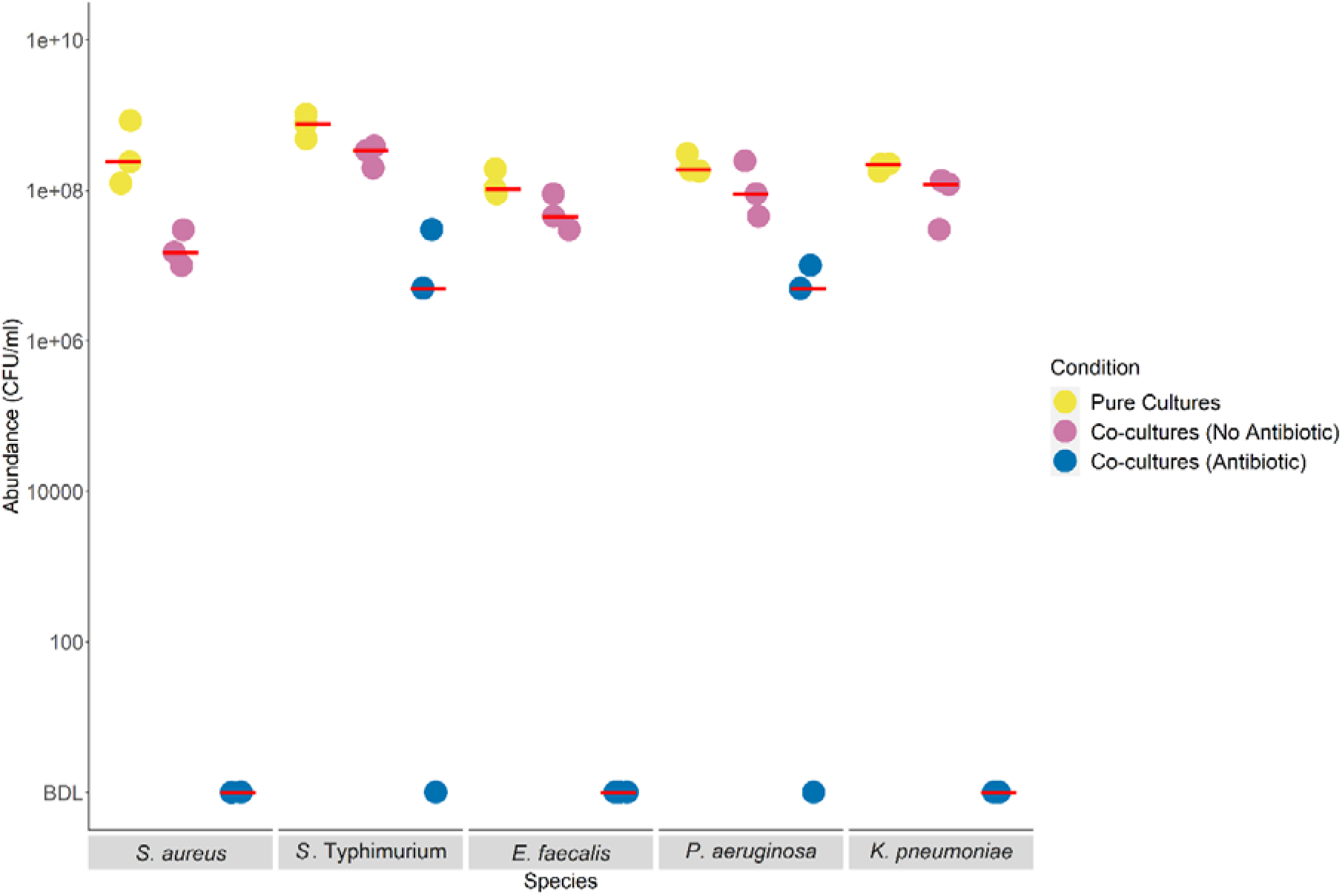
Only *P. aeruginosa* and *S*. Typhimurium show viable population growth in co-cultures with plasmid-carrying *E. coli* plus antibiotics. Abundance (CFU/ml) of each antibiotic-susceptible species excluding *E. coli* after 24h incubation is shown for three replicates in pure culture without antibiotics, in pairwise co-culture with the plasmid-carrying *E. coli* without antibiotics, or in pairwise co-culture with the plasmid-carrying *E. coli* with antibiotics. The red bars represent the median values for each condition. Note the detection limit in the antibiotic treatment here is quite high (5X10^6^ CFU/ml; below detection limit is signified as “BDL” on y-axis), due to the density of plasmid-carrying *E. coli* in those cultures.

## Discussion

In six-species communities, *E. coli* carrying a resistance plasmid detoxified the environment for other species. This had large effects on community structure, because some species profited more than others from detoxification. The identities of the species profiting most were explained by interspecies variation of their intrinsic population growth (measured in pure-culture) at degraded antibiotic concentrations (lower than the starting concentration). By contrast, two widely measured parameters in resistance evolution (MIC and pure-culture growth rate without antibiotics) only partially explained the success of *P. aeruginosa* and *S*. Typhimurium in detoxified community microcosms. Further experiments including pairwise co-cultures on agar, supernatant assays and extraneously changing the antibiotic concentration to mimic detoxification all supported the same pattern in terms of how detoxification changed community structure. We found no evidence that higher-order interactions among susceptible species influenced community-level responses to detoxification, despite evidence of these in some other scenarios (13,14). Thus, our results reveal community-level effects of detoxifying resistance mechanisms, including which species are most likely to increase in relative abundance upon antibiotic degradation.

The first important implication of our results is that the shift in community composition we observed upon plasmid-mediated detoxification was better explained by interspecies variation of the capacity to grow at degraded antibiotic concentrations than by classically measured parameters, including MIC and antibiotic-free growth rate. *P. aeruginosa* and *S*. Typhimurium grew better than other species at non-zero concentrations below the starting concentration, and were the only two species successfully exploiting detoxification in community treatments in our main experiment. These two species also had relatively high MICs, but so did *K. pneumoniae* (both in liquid and on agar), despite *K. pneumoniae* being unsuccessful in detoxified community microcosms. Thus, we do not argue that the MIC is uninformative, but it did not fully explain which species profited most in our experiments. Similarly, other species had similar growth rates in the absence of antibiotics compared with *P. aeruginosa* and *S*. Typhimurium, but did not thrive in detoxified community microcosms. This suggests a predictive understanding of community-level responses to antibiotics can benefit from additional information besides commonly measured parameters such as antibiotic-free growth rates and MICs (3,40). When there are detoxifying resistance mechanisms present, our results indicate responses of individual species, such as pathogens targeted by antibiotic treatment, depend critically on how they respond to degraded antibiotic concentrations (greater than zero but lower than that achieved at the beginning of treatment). In future work with greater numbers of species, interpretation of this type of interspecific variation could be facilitated by pharmacodynamic modelling, such as fitting Hill functions to dose response curves (41).

Our findings are likely to apply in other scenarios. First, the pOXA-48 plasmid is widespread and highly clinically relevant (42), so understanding this particular plasmid is important in itself. Second, the mechanism by which detoxification is encoded on this plasmid, via an antibiotic-degrading beta lactamase enzyme, is common among other plasmids, bacteria and resistance genes (3,8,10). Third, the species in our communities are all common in nature and associated with humans as commensals and/or pathogens. Moreover, the principle mechanism by which community composition changed in our experiment relies on interspecific variation of responses to antibiotic degradation, which we can expect to be pervasive also in communities with greater numbers of species. Fourth, we found similar patterns in terms of species-specific responses to detoxification both in liquid cultures and on agar, indicating this also applies in spatially structured conditions.

We were surprised to find no evidence of a role for horizontal transfer of the plasmid in community-level responses to detoxification. The pOXA-48 plasmid is generally expected to be horizontally transferrable (34,43). We do not rule out that there may have been some plasmid transfer in our experiments, but even if this occurred, it did not explain which species profited from detoxification (neither *P. aeruginosa* nor *S*. Typhimurium detectably acquired resistance). Our results are therefore relevant for scenarios where detoxifying resistance is encoded on non-mobile elements. In addition, we speculate detoxification itself may contribute to a reduced role for acquisition of resistance by susceptible species (via horizontal transfer or by chromosomal mutation) compared to scenarios where resistant species do not degrade the antibiotic. This is because detoxification should reduce the selective advantage of any newly arising resistant variants relative to their susceptible counterparts (we expect this advantage to be greatest above the MICs of susceptible strains but below the MIC of the resistant strain), resulting in a slower increase in transconjugant frequency. Nevertheless, future studies with other plasmids encoding both antibiotic-degrading and non-degrading resistance mechanisms and greater numbers of species would be informative.

In conclusion, we found carriage of a representative antibiotic-degrading resistance mechanism by one species strongly changed the responses of neighbouring species to antibiotic exposure in multispecies communities. This caused a shift in community composition and total abundance relative to equivalent communities without resistance. Surprisingly, the extent to which susceptible species benefited from antibiotic degradation depended on their intrinsic ability to grow at degraded antibiotic concentrations, rather than on horizontal gene transfer, higher-order interactions among susceptible species, or classical parameters such as the MIC. This improves our ability to predict how individual species and whole communities respond to antibiotics. Our data show responses of individual species depend not only on their own susceptibility to antibiotics, but which resistance mechanisms are circulating in their neighbours. Therefore, as clinical metagenomic data becomes more widely available and applicable in treatment contexts and diagnostics (44), annotation of potentially degrading resistance mechanisms such as beta-lactamases in microbiome samples, even if they are not detected in targeted pathogens, may improve our ability to predict treatment outcomes. The relevance of this is further supported by evidence that antibiotic-degrading mechanisms, like that in our experiments, are common in human-associated communities (45).

## Acknowledgements

We thank Adrian Egli at Universitätsspital Basel for clinical plasmid-carrying strains. We thank Steve Kett and Justus Fink for the feedback on the manuscript. We thank Helena Klein for advice on visualisation. Funding source: Swiss National Science Foundation project number 310030_192428.

