## Supplementary Material for "Antibiotic-degrading resistance changes bacterial community structure via species-specific responses"

### Contents

Table S1: Description of bacterial strains

Figure S1: Conjugative transfer of pOXA-48 plasmid from native clinical isolate to K-12 MG1655

Figure S2: Growth dynamics of bacterial strains

Figure S3: Species-specific benefits of detoxification on agar. Images of antibiotic exposure protection of susceptible strains on agar.

Figure S4: Images of antibiotic exposure protection of assembled communities on agar.

**Table S1.** Description of all the strains used in this study. ATCC refers to American Type Culture Collection.

| Species | Strain | Citation or ATCC reference |
| --- | --- | --- |
| *Escherichia coli* | K-12 MG1655 (CmR, Δ*galK::cat*) without pOXA-48 plasmid | (1) |
| *Escherichia coli* | K-12 MG1655 (CmR, Δ*galK::cat*) with pOXA-48 plasmid |  |
| *Staphylococcus aureus* | Type Strain | ATCC 12600 |
| *Salmonella enterica* serovar Typhimurium | SL1344 | ATCC SL1344 |
| *Enterococcus faecalis* | JH2-2 | (2) |
| *Pseudomonas aeruginosa* | PAO1 | (3,4) |
| *Klebsiella pneumoniae* subsp. *pneumoniae* | Type Strain | ATCC 13883 |

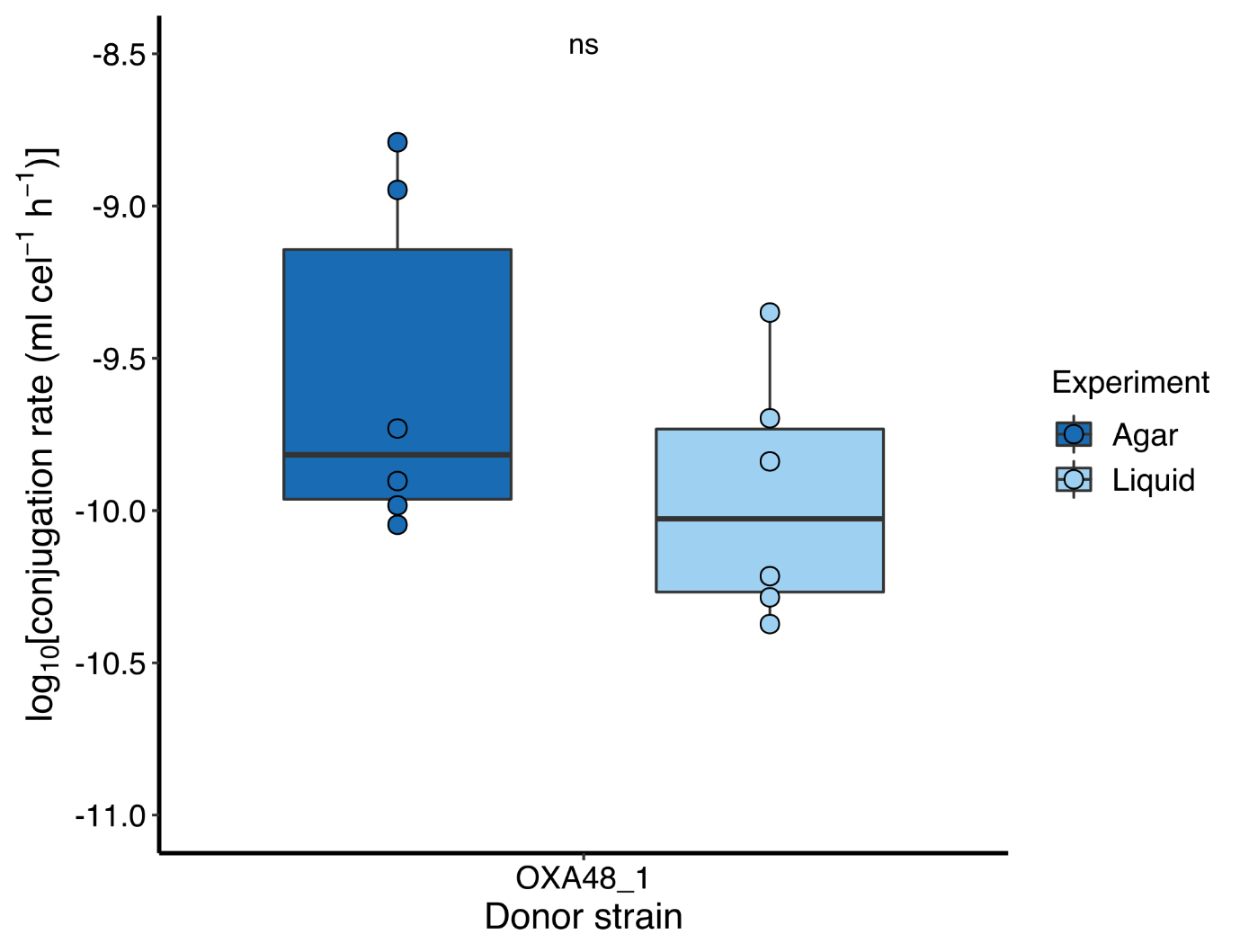
**Figure S1: Conjugation rate of** **pOXA48 into a chloramphenicol resistant *E. coli* K-12 MG1655 (CmR, Δ*galK::cat*) on agar and in liquid** (5,6). For the agar assay, we first grew six colonies each of the *E. coli* clinical strain (native plasmid host) and the recipient (MG1655_CmR) in 2 ml of LB. After 3.5 h at 37 °C with 180 r.p.m., we pelleted each culture (1 ml centrifuged 5 min at 1500G), resuspended in 100 µl LB, then mixed donor:recipient cultures 1:1 (*v*:*v*) and spotted onto LB agar plates. After 1 h at 37°C, spots were resuspended in sterile NaCl 0.9% and appropriate dilutions plated on selective plates (ampicillin 100 mg l^−1^, chloramphenicol 50 mg l^−1^ and a combination of both). For the liquid assay, after mixing the independent overnight cultures of the donor and recipient in a 1:1 ratio, we incubated a 1000-fold dilution for 24 h at 37°C. After 24h, appropriate dilutions were plated on selective plates. Plasmid presence was verified by *bla*_OXA-48_ gene amplification by PCR (Fw: GGCGTAGTTGTGCTCTGGAA; Rv: CCAACCGACCCACCAGCCAA). Conjugation rates were determined using the end-point method (7).

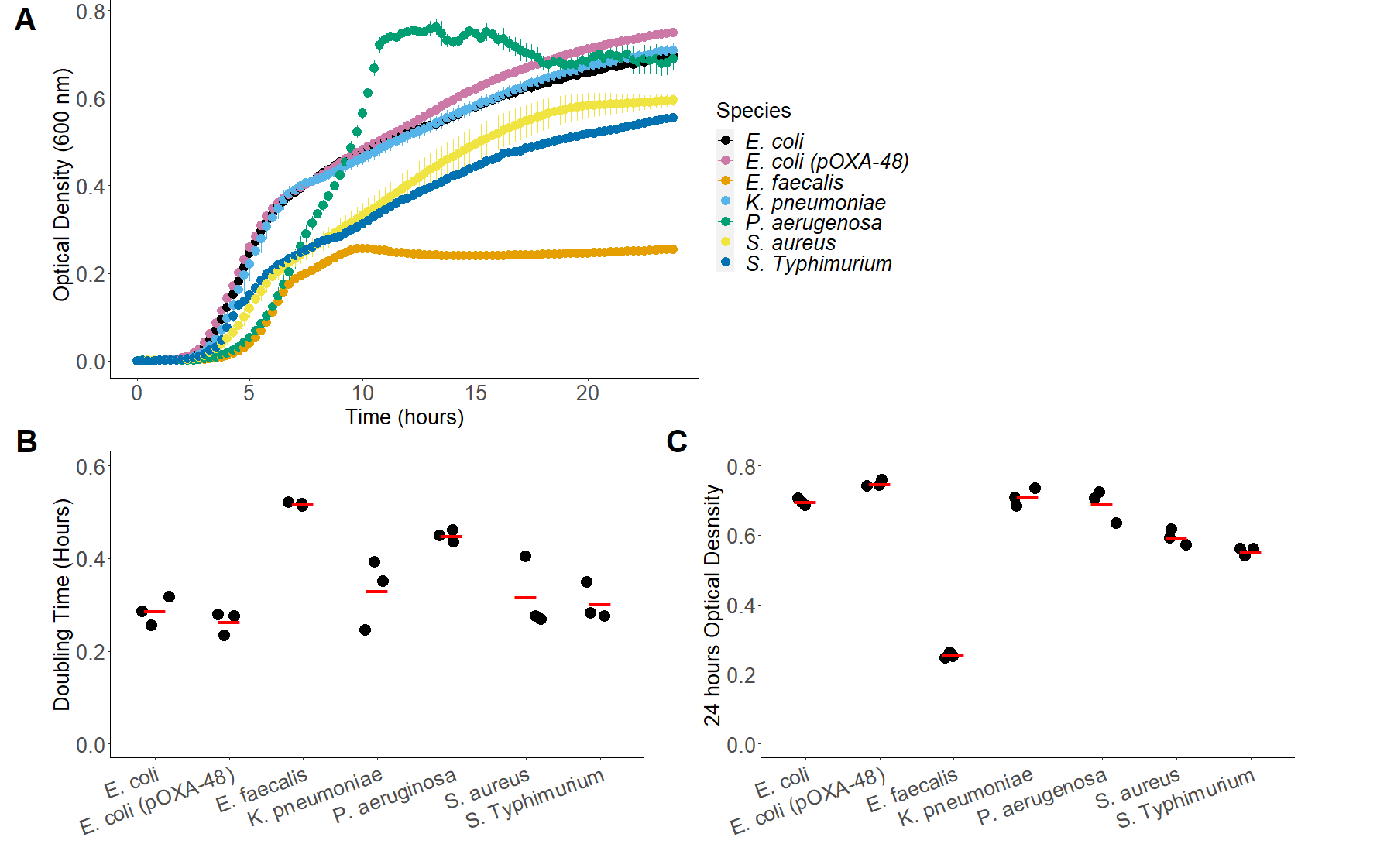

**Figure S2**. Growth dynamics of all strains over 24 hours. **(A)** Abundance of each strain (see legend) measured by optical density over 24h. Points show means ±95% confidence intervals, with three replicates per strain. **(B)** Growth rates estimated from the data shown in panel A, estimated using a sliding-window approach (see Methods). Average doubling time varied among strains (p<0.05 by one-way ANOVA); post-hoc Tukey’s HSD showed *E. faecalis* had a doubling time equivalent to that of *P. aeruginosa* and higher than all other strains, whereas *P. aeruginosa* had a doubling time equivalent to that of *E. faecalis* and *K. pneumoniae* and higher than all other strains. **(C)** Final abundances, inferred from final OD measurements after 24 hours for each species in panel A. Final abundance varied among species (p<0.05 by one-way ANOVA); post-hoc Tukey’s HSD indicated *E. faecalis* had a lower average abundance than all other strains, *S. aureus* and *S*. Typhimurium were similar, significantly higher than *E. faecalis* and significantly lower than all other strains.

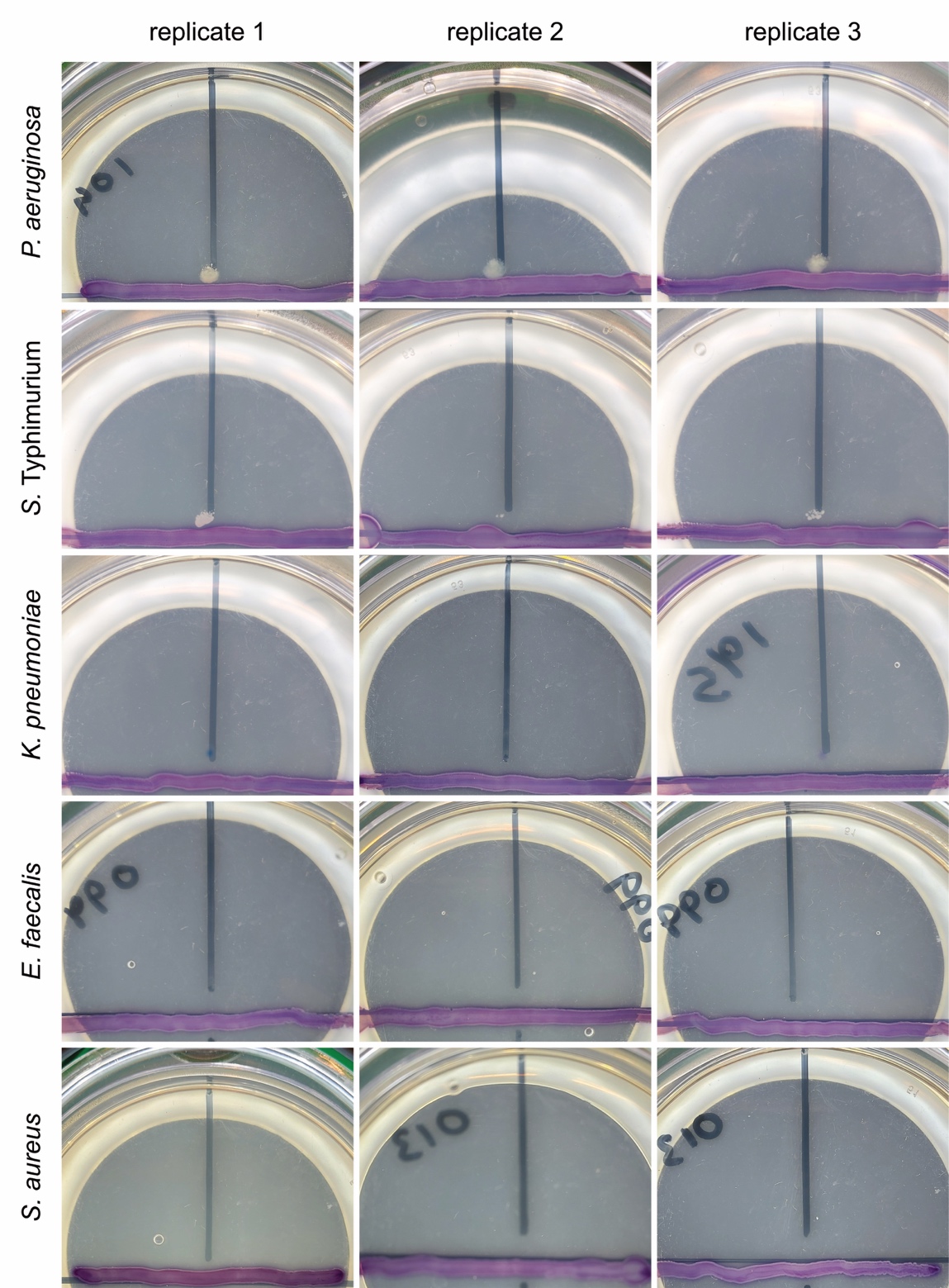

**Figure S3.** Two out of five antibiotic-susceptible species (*P. aeruginosa* and *S*. Typhimurium) show visible growth on antibiotic-agar in proximity to a resistant, plasmid-carrying *E. coli* strain. Each susceptible species (rows) was streaked on chromatic agar with antibiotics (down the vertical black line) perpendicular to the pOXA-48 carrying *E. coli* (the purple horizontal line of bacterial growth) and incubated for 24 hours. Each combination was replicated three times (the three columns in each row). Three out of three replicates for *P. aeruginosa* and two out of three for *S*. Typhimurium were qualitatively identical.

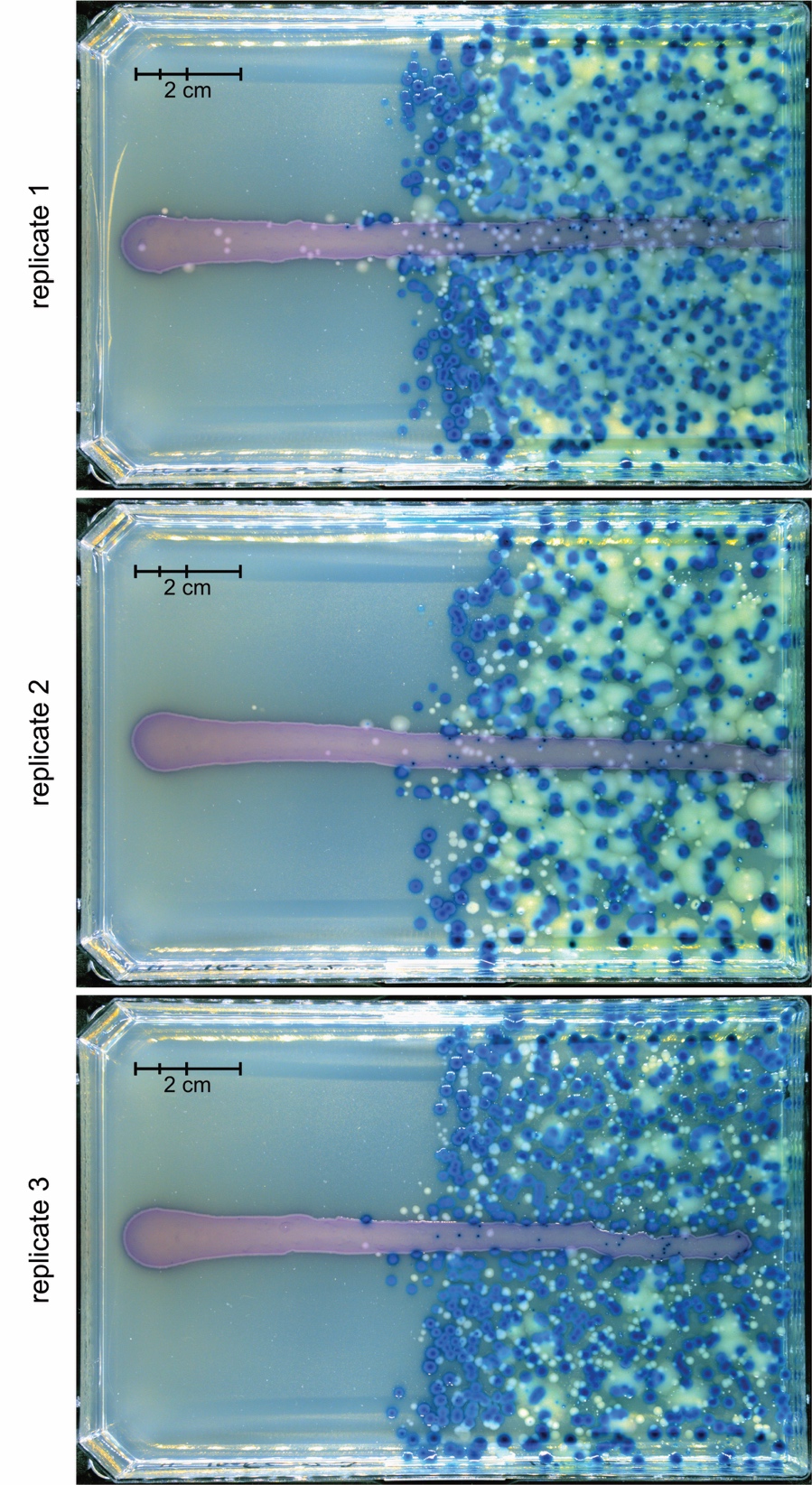

**Figure S4.** Multispecies communities plated on chromatic agar with a gradient of piperacillin+tazobactam (left-to-right; maximum concentration in overlay agar = 5 µg/ml piperacillin and 1 µg/ml tazobactam) with the plasmid-carrying *E. coli* strain streaked horizontally (purple). *S.* Typhimurium appears as defined, small off-white colonies, *P. aerugionosa* appears as diffuse off-white colonies, *K. pneumoniae* appears dark blue, *E. faecalis* appears light blue and *S. aureus* appears as small white colonies (see also Fig. 3B for labelling).
